# Employing active learning in the optimization of culture medium for mammalian cells

**DOI:** 10.1101/2022.12.24.521878

**Authors:** Takamasa Hashizume, Yuki Ozawa, Bei-Wen Ying

## Abstract

Medium optimization is a crucial step of cell culture for biopharmaceutics and regeneration medicine. It remains challenging, as both media and cells are highly complex systems. To address the issue, we tried active learning to fine-tune the culture medium by combining the high-throughput assay and machine learning. As a pilot study, the cell line HeLa-S3 and the gradient-boosting decision tree algorithm were used. The regular and time-saving approaches were developed, and both successfully fine-tuned 29 components to achieve improved cell culture than the original medium. The fine-tuned media showed a significant decrease in fetal bovine serum and the differentiation in vitamins and amino acids. Unexpectedly, the medium optimization raised the cellular NAD(P)H abundance but not the cell concentration owing to the conventional method used for cell culture assay. Our study demonstrated the efficiency of active learning for medium optimization and provided valuable hints for employing machine learning in cell culture.

## Introduction

Efficient approaches for medium optimization of cell culture were highly required. Various methods were developed, as mammalian cells were commonly used in the medical and biopharmaceutical fields^1^. Besides the development of the cell lines^2, 3, 4,^ the culture media were intensively studied to improve the performance of developed cell lines^5^. The composition of the culture medium, e.g., sugars, amino acids, fatty acids, vitamins, trace elements, and salts^6^, were usually required to be fine-tuned for cell growth and production^7, 8, 9^. As the contribution of the medium components to the cellular metabolism was complicated^10^, the medium optimization remained challenging. The conventional method OFAT (One-Factor-At-Time) fine-tuned the medium components one by one ^11^. The statistical method DOE (design of experiments) was efficient when the medium components requiring adjustment were fewer than ten^12^. The mathematical method RSM (response surface methodology)^13, 14^ used the quadratic polynomial approximation, which was too simple to represent the comprehensive interaction between medium and cell^15^. The approach of machine learning (ML) was proposed to overcome these limitations. It was tried for medium development^16, 17^ and showed higher performance than DOE and RSM^15, 18^.

Active learning was recently proposed to improve the performance of ML-assisted medium optimization. The efficiency of medium optimization largely depended on the prediction accuracy of the ML model itself. Active learning was expected to improve prediction accuracy with a small number of datasets by allowing the ML models to select data for training^19, 20^. It was reported to be practical in drug discovery^21, 22, 23^, as well as successfully optimized the buffer composition for protein biosynthesis in a cell-free system ^24^. So far, applying active learning for medium optimization of mammalian cell culture is still under investigation. For instance, of the studies on medium development for CHO cells in the past 20 years, 37% considered only one additive, 37% used OFAT, 15% used DOE, and none used machine learning or active learning^25^. Medium optimization of other cell lines used the ML algorithms without active learning^16, 26^. Whether active learning was applicable to optimize the culture medium of mammalian cells was an intriguing issue.

The present study first tried active learning combining explanatory ML with experimental validation to optimize the medium composition for improved cell culture^25,27^. Although the medium components all influenced cell growth and production, the contributions of medium components to cell culture focused on amino acids, which were differentiated in cell viability, growth, and bio-production^28, 29^. Whether and how the other medium components contributed to the cell culture remained unclear, as the ML models were usually black boxes. The ML algorithm, gradient-boosting decision tree (GBDT), was employed as it was a white box. Our previous study used GBDT to explore the contribution of medium components to bacterial growth and successfully observed the chemical dependence of different components as the survival strategy^30^. Applying GBDT in active learning for medium optimization of mammalian cells could allow us to identify the contribution of individual medium components to cell culture.

## Results

### Experimental design for acquiring the training data

The cell line HeLa-S3 was used in the study, as it could grow in a free-floating mode, making it easy to evaluate. The initial cell concentration was determined at 10^4^ cells/ml, as higher and lower concentrations led to a reduced growth rate and extended culturing time (Fig. S1). Quantitative evaluation of the cell culture was decided by testing multiple methods, i.e., counting the cell particles, cell imaging analysis, chemical reaction assay, and cell stain and counting (Fig. S2). Considering the time consumption of operation and the measurement range of cell concentration, the chemical reaction assay upon the cellular NAD(P)H abundance was selected, where the cell concentration was evaluated as the absorbance at 450 nm (A450). This method was efficient and convenient for acquiring an extensive data set for ML, as it could be performed in a high-throughput manner. In addition, the medium components subjected to optimization were determined according to the composition of the commonly used Eagle’s minimum essential medium (EMEM), which comprised approximately 31 components. Except for phenol red and penicillin-streptomycin, 29 components were used to prepare the medium combinations for active learning (Fig. 1A). The concentration gradients of these components were varied on a logarithmic scale to acquire a broad data variation. Finally, cell culture in 232 medium combinations was performed, and the temporal changes of cell culture were measured at 24 or 48-h intervals (Fig. 1B). Biological replication (N=3∼4) was conducted for each sampling point, resulting in thousands of A450 records. Note that there’s no manual (personal) bias in preparing the medium combinations, as the commercially purchased EMEM and the lab-made EMEM showed equivalent effects on cell culture (Fig. S3)

**Figure 1.**
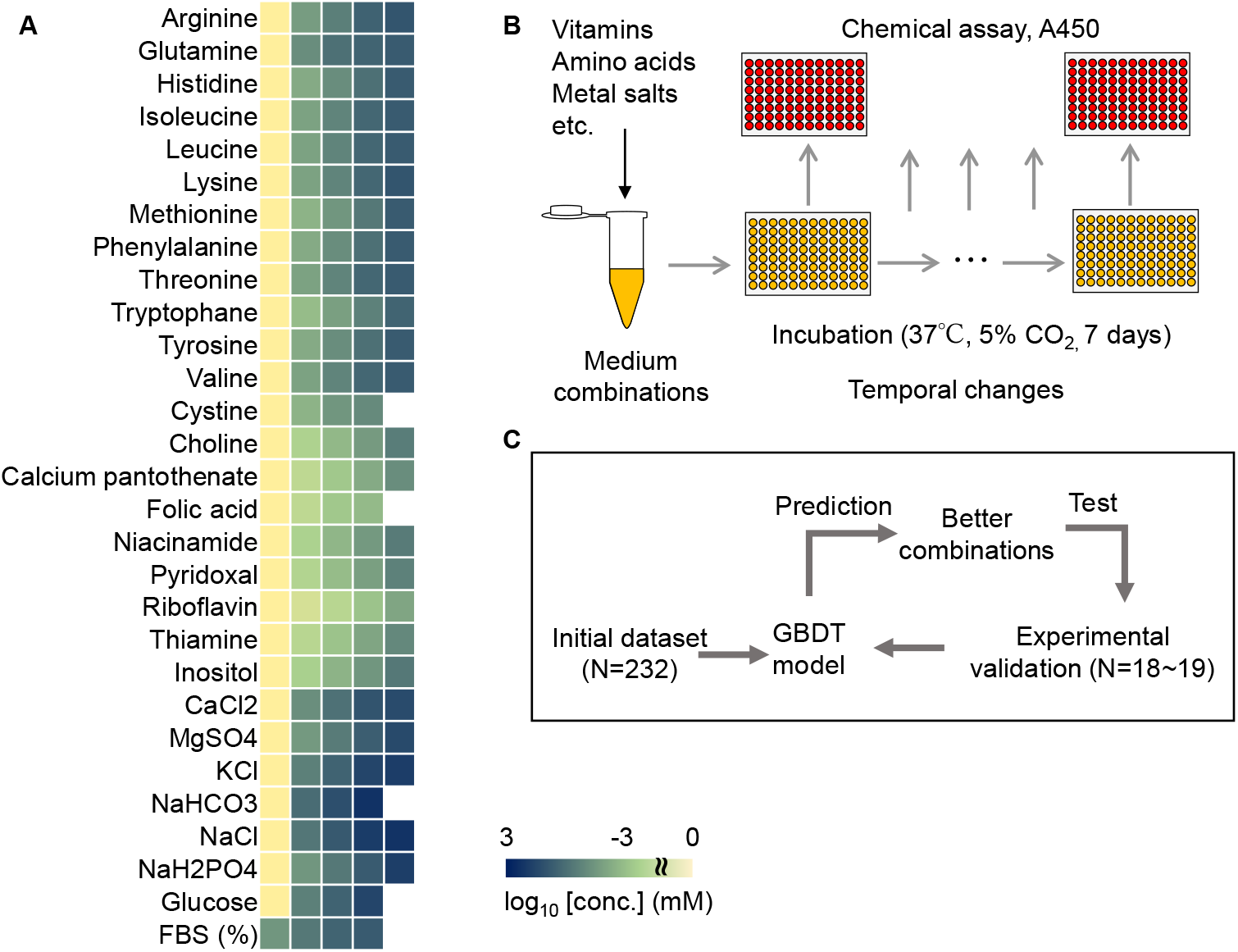
Experimental data acquisition and active learning. **A**. Medium components and their concentrations used in the initial dataset for active learning. Color gradation indicates the concentration gradients shown on a logarithmic scale. **B**. Schematic drawing of experimental procedure for experimental data acquisition. **C**. Flowchart of the active learning for medium optimization.

### Regular and time-saving modes of active learning for medium optimization

Active learning was performed to search for the medium combination of improved cell culture. As a regular mode of active learning, A450 of the cell culture at 168 h was used for the training because it was roughly the time point reaching the saturated cell concentration (Fig. S1). The GBDT model was used to predict the medium combinations of better cell culture, i.e., higher A450, and the learning loop was started with the dataset comprising 232 medium combinations (Fig. 1C). Every 18∼19 predicted combinations were subjected to experimental validation. The experimental results were added to the training data. The procedure of model building, medium prediction, experimental validation, and training (Fig. 1C) was repeated for four rounds (Fig. 2A). Resultantly, both the A450 values of the cell culture and the accuracy of the GBDT models were elevated. The cell culture was significantly improved in Round 3 and remained comparable after Round 4 (Fig. 2A). We assumed that either the methodology or the cell culture reached its limitation. In addition, the prediction accuracy was improved gradually along with the rounds exceeded (Fig. 2B, Fig. S4), indicating that the active learning finetuned the medium combination associated with increasing ML accuracy.

**Figure 2.**
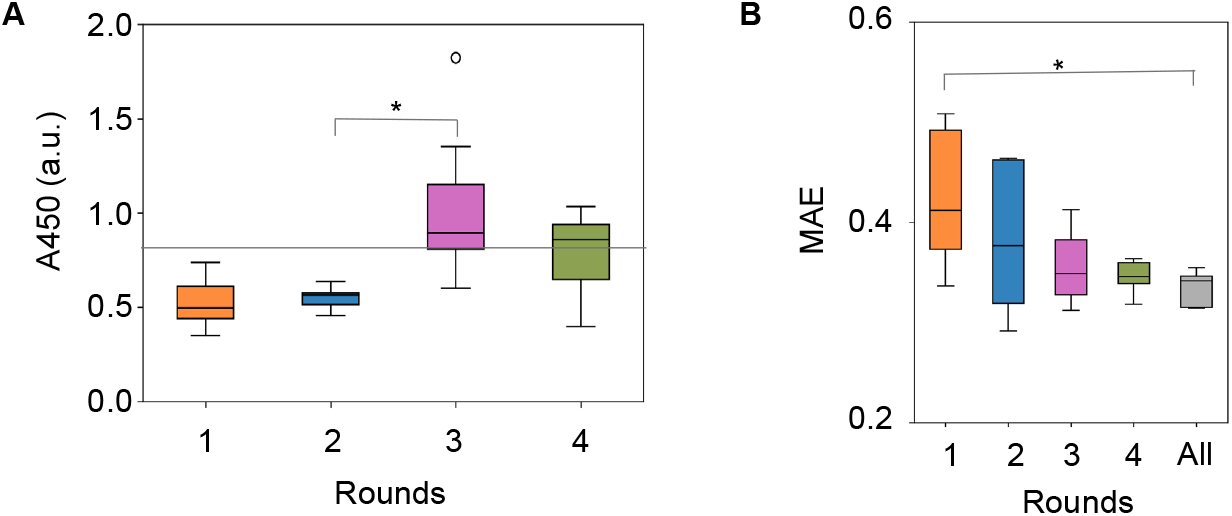
Regular mode of active learning for medium optimization. **A**. Boxplots of A450 of the cell culture at 168 h. Every 18 to 19 medium combinations predicted by the GBDT model using the culture data at 168 h are tested. The gray horizontal line indicates the cell culture with EMEM. **B**. Boxplots of the prediction accuracy of the GBDT model built in each round. MAE was calculated by 5-fold nested cross-validation. “All” indicates the accuracy evaluated using the entire dataset from the initial to Round 4. Asterisks indicate statistical significance by Mann-Whitney’s U test (p<0.05).

Subsequently, whether the active learning loop could be shortened was tested. As the data acquisition step (168 h) was time consumed, whether A450 of the cell culture at an earlier time could be used for predicting the cell culture at 168 h was investigated. As a time-saving mode, A450 of the cell culture at 96 h was employed for active learning because a highly significant correlation of A450 was detected between 96 h and 168 h (Fig. S5). The 232 medium combinations associated with the A450 values of the cell culture at 96 h were employed as the initial dataset for active learning (Fig. 1C). Both the A450 of the cell culture at 96 h and the prediction accuracy were improved (Fig. 3A-B, Fig. S4), as observed in the regular mode. Intriguingly, A450 of the cell culture at 168 h was significantly increased (Fig. 3C), despite the ML using the culture data acquired at 96 h. The medium combinations for improved cell culture were successfully achieved using the culture data earlier than required, which shortened hundreds of hours for medium optimization in the present case (Fig. S6). It demonstrated that the time-saving mode was practical for ML-assisted medium optimization for improved cell culture.

**Figure 3.**
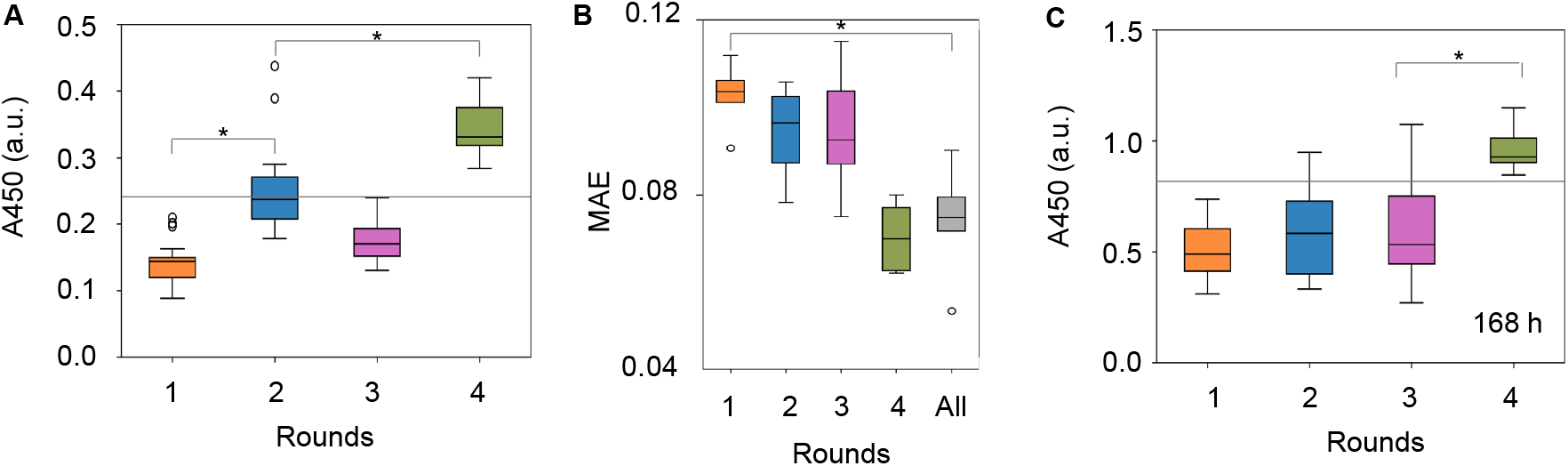
Active learning for medium optimization in time-saving mode. **A**. Boxplot of A450 of the cell culture at 96 h. Every 18 to 19 medium combinations predicted by the GBDT model using the culture data at 96 h are tested. **B**. Boxplot of the prediction accuracy of the GBDT model built in each round. MAE was calculated by 5-fold nested cross-validation. “All” indicates the accuracy evaluated using the entire dataset from the initial to Round 4. **C**. Boxplot of A450 of the cell culture at 168 h. Every 18 to 19 medium combinations predicted by the GBDT model using the culture data at 96 h are tested. The gray horizontal lines indicate the cell culture at 96 or 168 h with EMEM. Asterisks indicate statistical significance by the Mann-Whitney U test (p<0.05).

### Contribution and composition of fine-tuned medium components

The contribution of medium components to cell culture was estimated by GBDT. The entire datasets acquired through active learning, i.e., 302 and 403 medium combinations in the regular and time-saving modes, respectively, were used (Fig. S7, Fig. S8). The importance of each component in the two modes was estimated. The top ten components that mainly contributed to cell culture partially overlapped (Fig. 4A, B). It indicated both the regular and the time-saving modes fine-turned the similar components, e.g., metal salts and FBS, for improved cell culture. The finding partially explained why the medium optimization at 96 h improved cell culture at 168 h. Intriguingly, NaCl and CaCl_2_, but not FBS, were the primary components deciding the cell culture in the regular and time-saving modes, respectively. Although FBS usually contained calcium ions^31^ and 1∼3 mM CaCl_2_ was generally supplied in the cell culture ^6^, adjusting the concentration of calcium ions was essential for cell growth because either excess or deficient calcium ions would induce cell apoptosis^32, 33, 34, 35^. In addition, high osmotic pressure, caused by the high concentration of NaCl in the medium, would arrest cell growth^36^. The results indicated that the cellular permeability, regulated by NaCl and CaCl_2_, rather than the growth factors provided by FBS, might present the highest priority in cell culture.

**Figure 4.**
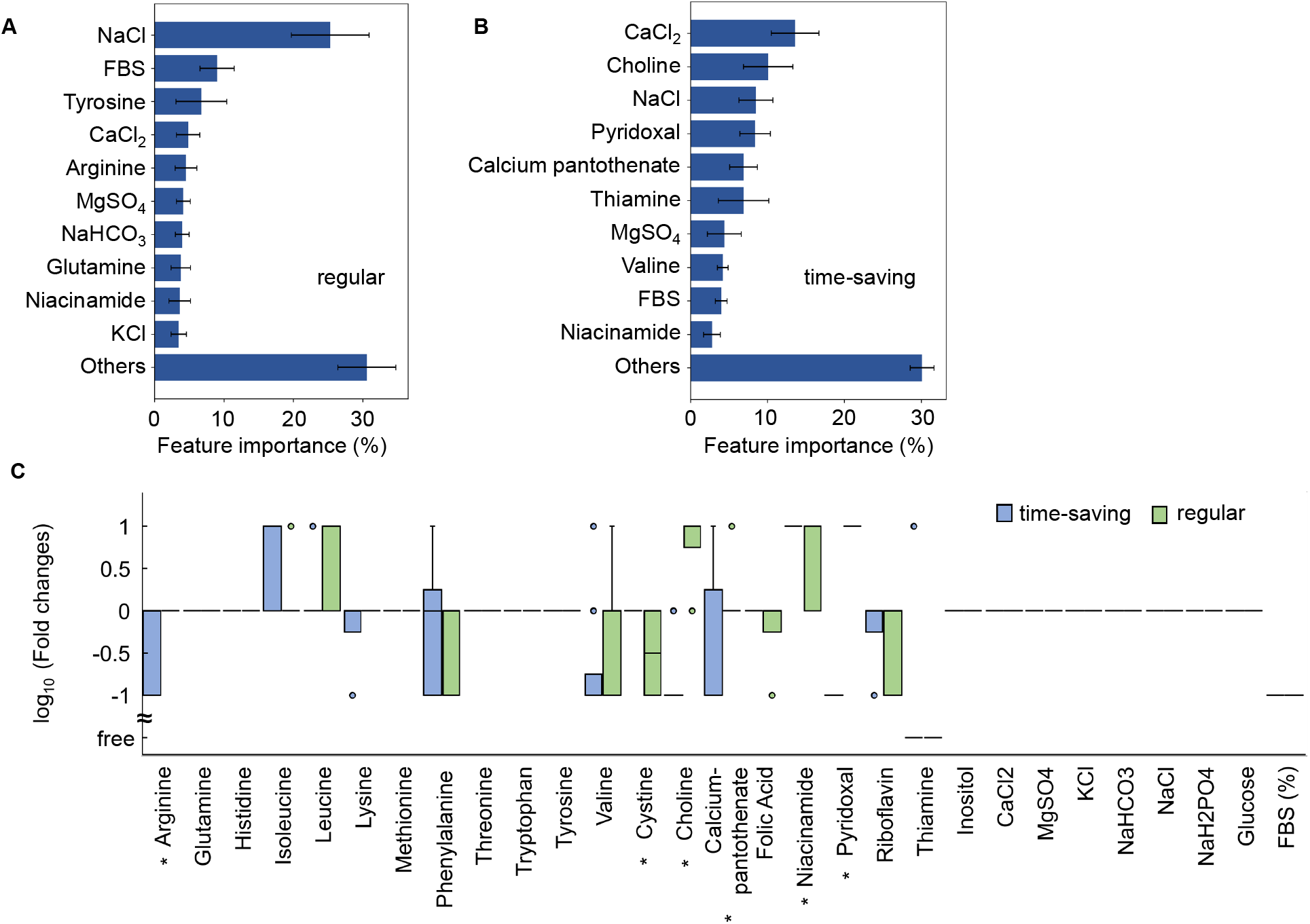
Contribution and composition of the optimized media. **A**. Relative contribution of medium components to cell culture at 168 h. The top ten components and the sum of the remaining 19 components, i.e., Others, are indicated. Five-fold cross-validation of GBDT in the regular mode was applied. Standard deviations of five replications are indicated. **B**. Relative contribution of medium components to cell culture at 96 h. The top ten components and the sum of the remaining 19 components, i.e., Others, are indicated. Five-fold cross-validation of GBDT in the time-saving mode was applied. Standard deviations of five replications are indicated. **C**. Fold changes in concentrations of medium components. Concentrations of the 29 components in the top ten optimized media were compared to those in EMEM. Green and blue represent the media predicted with regular and time-saving modes. Fold changes in concentration are shown in the logarithmic scale. Asterisks indicate statistical significance by Mann-Whitney U-test (p<0.05).

The compositions of the best ten media predicted with the regular and time-saving modes were compared. The mean concentration of the component presented in the ten best media was compared with its concentration in EMEM. Most components presented similar concentrations predicted with the two modes. However, six out of 29 components showed significant differentiation in concentrations, either amino acids or vitamins (Fig. 4C). It indicated that the various combinations of amino acids and vitamins could improve cell culture. Additionally, in all 20 media, FBS was as low as 10% of the commonly used abundance. FBS contained a variety of factors essential for cell growth, including trace elements (vitamins and minerals), hormones, free radical scavengers, mitogenic growth factors, and so on^37^, which were absent in the basal media^38^. Generally, 10∼20% of FBS in the media was considered suitable for cell culture, as reported in enterocytes^39^, stem cells^40^, and hybridoma^41^. Such a reduced amount of FBS was supposed to be a preferred trend, considering the risk of contamination, the cost, and the batch-to-batch variability in cell production^42, 43^.

### Comparison of the fine-tuned media to the original medium

The optimized media showed higher A450 than the original medium EMEM, regardless of the regular and time-saving modes. The best ten medium combinations (i.e., showed the highest A450) predicted with the regular and time-saving modes were prepared for the experimental verification of the cell culture at 168 h. The predicted media showed better performance, i.e., higher A450, than the original medium EMEM (Fig. 5A). It demonstrated that both modes resulted in successful medium optimization, although the media predicted with the regular mode showed better performance than those predicted with the time-saving mode.

**Figure 5.**
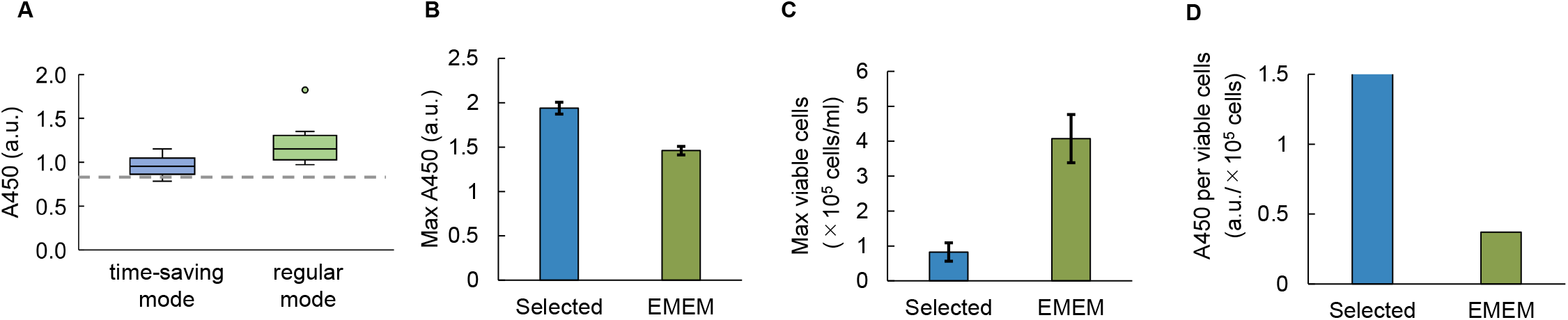
Performance of the ML-optimized media. **A**. Comparison of the optimized media predicted with the regular and time-saving modes. The top ten media, which showed the highest A450 absorbance of cell culture, are shown. The broken line indicates the cell culture in EMEM. **B**. Comparison of A450 between the selected and EMEM media. Standard errors are indicated. **C**. Comparison of the number of viable cells between the selected and EMEM media. Standard errors are indicated. **D**. Comparison of cellular A450 between the selected and EMEM media. The ratio between the mean A450 and the mean number of viable cells is shown.

In addition, a direct comparison of the optimized medium to the original medium was performed. The best of the 20 predicted media was selected to compare with the commercially purchased EMEM. The cell culture was performed in a 1-mL volume as a scale-up compared to the culture volume used in active learning. Intriguingly, the selected medium showed higher A450 (Fig. 5B) but a lower number of viable cells than EMEM (Fig. 5C). It seemed that the increased A450 per cell caused the improved A450 in the selected medium (Fig. 5D). Active learning of medium optimization must have raised the cellular abundance of NAD(P)H rather than the number of cells. This finding made us reconsider whether the commonly used chemical assay of cell culture indeed reported the cell concentration solely. The methodology used for experimental data collection played a crucial role in the final output of the fine-tuned parameters.

## Discussion

Cell culturing time was one of the time-consuming processes in medium development. As known, the mammalian cell culture requires a longer time than the *in vitro* reaction and bacterial culture. Quick optimization of medium composition was often performed by statistically reducing the experimental test conditions^44^. To acquire the experimental dataset for ML-assisted medium optimization, shortening the data acquisition time might be an alternative approach for quick optimization. The present study made a first trial of reducing the experimental time (Fig. S6). The information on the early phase culture successfully predicted the output of the late phase culture (Fig. 3C). It indicated that the time-saving active learning was practical in accelerating the medium development for regeneration medicine and the R&D of biopharmaceuticals, which was highly required^45^. In the present study, active learning of medium optimization successfully, somehow unexpectedly, raised the cellular NAD(P)H abundance but not the cell concentration. The NAD(P)H abundance was known to be proportional to the number of viable cells^46^, which relied on the thinking that the NAD(P)H abundance per viable cell was constant. However, the results showed that the increased NAD(P)H abundance did not always cause an increased number of vital cells (Fig. 5D). Cellular NAD(P)H abundance could be raised by medium optimization. We assumed that the fine-tuned media benefited the metabolism for increased NAD(P)H abundance in cells. The upper limit of cellular NAD(P)H abundance might be the reason for the failure of active learning to increase A450 after Round 4 (Fig. 2). In an alternative viewpoint, the cellular capacity of NAD(P)H could be addressed by active learning. This finding brought us a novel idea of using active learning to search for the biological limitation of living cells, e.g., the highest growth rate.

Why the optimized medium increased the cellular NAD(P)H might be associated with glucose metabolism, medium pH, and reactive oxygen species (ROS) effects. Firstly, the chemical assay using water-soluble tetrazolium (WST) depended on both the viable cells and the glucose metabolism^47^. Enhanced glucose metabolism without changes in cell concentration could raise the absorbance caused by WST. As the optimized media and EMEM contained the equivalent amount of glucose, the changes in glucose metabolism might not be the primary cause. Secondly, the chemical assay could be affected by the pH of the medium^48^. As the optimized media remained at the same pH as EMEM, the absorbance bias due to pH could be excluded. Finally, the increased NAD(P)H abundance could be caused by oxidative stress. Cancer cells protected themselves from ROS by increasing NAD(P)H^49, 50^, which converted oxidized glutathione to reduced glutathione^51^. ROS might be attributed to the medium components^52^, such as FBS and thiamine. FBS comprises various enzymes and macromolecules as antioxidants, and thiamine acts as an antioxidant^53, 54^. Depletion of the antioxidants could trigger oxidative stress^55, 56^. The optimized media comprised ten-fold fewer FBS than EMEM and no thiamine (Fig. 4C), which probably led to the increased oxidative stress indicated by the cellular NAD(P)H abundance.

Vitamins and amino acids were highly prioritized in the time-saving and regular modes, indicating the temporal differentiation in chemical function. In the time-saving mode, five of the ten determinant components were vitamins (Fig. 4B), closely related to the cellular NAD(P)H abundance mediated by ROS. The choline pathway synthesized serine and glycine, which were required for antioxidant glutathione synthesis^57^, were absent in the media. Pyridoxal inhibited the formation of superoxide radicals^58^. Pantothenic acid prevented ROS-induced apoptosis^59^. Niacinamide was the amide of niacin, a precursor of NAD^+^ and NADP^+53^. The high priority of these vitamins indicated they played a crucial role in deciding the early phase of cell culture. In the regular mode, tyrosine, arginine, and glutamine were the determinants. The abundance of amino acids was related to cell viability^60^, as the excess amount could accumulate toxic metabolites^61^. Metabolisms of the three amino acids produced ammonia, 4-Hydroxyphenylpyruvate, and dimethylarginine^62^, which were toxic to cells^63, 64^ and caused growth inhibition^65^ or cell apoptosis^66, 67^. The high priority of these amino acids suggested that an adequate amount was essential to maintain the balanced metabolism for cell viability in the late phase of cell culture.

The study demonstrated active learning was effective in medium optimization and the time-saving mode was practical. However, additional “wet” and “dry” issues should be considered to improve the accuracy of ML, i.e., the evaporation during cell culture and the data selection. Firstly, water evaporation in a CO_2_ incubator during incubation was a critical issue to consider. The present study observed ∼ 13.3% evaporation with a locational variation (Fig. S9). The changes in cell concentration might be biased by the degree of water evaporation that altered the culture volume. Such bias reduced the precision of the experimental data, consequently affecting the quality of the training set. Considering water evaporation as one of the features in the ML models might be a solution.

Secondly, uncertainty sampling was commonly employed to improve the ML models in active learning^20^. In the present study, uncertainty sampling was absent in the datasets, as only the medium combinations for better cell culture were selected for experimental validation in active learning. Additional experimental data of worse cell culture, as uncertainty sampling, for training was required to improve the accuracy of ML models for further development of culture medium.

## Materials and Methods

### Cell line and culture

HeLa-S3 cells were from the RIKEN Cell Bank (Tsukuba, Japan). Cell culture was performed at 37°C in a CO_2_ incubator (E-50, As One) supplied with 5% CO_2_^68^. Cells were cultured in the multiwell plates (Iwaki) with a culture volume of 0.5 or 1 mL in 48-or 24-well microplates, respectively. Multiple wells were used for biological replications. The cultured cells were detached with PBS (-) (Wako) supplemented with trypsin-EDTA (Sigma). The cells were collected by centrifugation and re-suspended in the cryopreservation solution (Wako) with trypan blue (Wako). The cells were counted in a hemocytometer (DHC-N01, Nano Entek) with a microscope (ECLIPSE Ts2, Nikon)^69^. The number of viable cells was evaluated accordingly.

### Preparation of cell stocks

Cell stocks for repeated cultures were prepared to reduce experimental errors^70^. The cells stored in liquid nitrogen were thawed at 37°C and suspended with 4 ml of Eagle’s Minimum Essential Medium (EMEM, Wako) for medium exchange. Subsequently, the cells were collected and suspended in 10 mL of EMEM, supplemented with 1% penicillin-streptomycin solution (Wako) and 10% FBS (Japan Bio Serum). The cells were cultured in 10 cm dishes (Violamo) for two days at 37°C in a CO_2_ incubator and were counted in the hemocytometer, as described above. The cell culture was divided into 1 mL and dispensed into 1.2 ml cryotubes (Biosigma), frozen at -80°C for 24 h, and finally stored in liquid nitrogen for future use. Resultantly, dozens of identical cell stocks were prepared.

### Preparation of medium combinations

the medium combinations were prepared with 31 commercially available compounds, in which choline chloride and pyridoxal hydrochloride were from Tokyo Chemical Industry, FBS from Japan Bio Serum, and the remaining were from Wako. The abundance of penicillin-streptomycin and phenol red was maintained at 1% and 0.03 mM, respectively. The concentrations of FBS were changed in the range of 0.1 to 10 %. The remaining 28 components were changed from zero to 10∼100 folds of their concentrations in EMEM. In brief, four to five different concentrations were used for each component, and the changes were roughly on a logarithmic scale, as described previously^30^. The medium combinations were prepared by mixing the stock solutions of individual components, prepared at high concentrations in advance, as described previously^30^. The stock solutions were sterilized by the sterile syringe filters (Merck) with hydrophilic PVDF membranes of 0.22 μm pore size and were dispensed in a small volume and stored at -20°C for future use. Note that all the medium combinations were prepared immediately before use, and the stock solutions were used for a single time.

### Cell counting according to single-particle analysis

A particle size analyzer (Multisizer 4, Beckman Coulter) was used to count the number of cells. The cell culture in a 48-well microplate (Iwaki) was suspended in 10 ml of ISOTON II (Beckman Coulter). Every 100 to 500 μl of the suspended solutions were flowed for particle analysis, according to the manufacturer’s induction. The size of the particles within the range from 8 to 12 μm in diameter was gated as the cells. The mean value of the biological replication was calculated as the cell concentration.

### Cell counting by imaging analysis

The cells cultured in a 48-well microplate (Iwaki) were imaged with BioStudio-T (Nikon) according to the manufacturer’s induction. The image of the cell culture was analyzed with the software NIS-Elements (Nikon), and the number of cells was evaluated automatically.

### Temporal changes of cell culture evaluated by chemical assay

The cell culture was performed in 200 μl with the 96-well microplates (Iwaki), and the culture condition was described above. As explained elsewhere^71^, only the wells in the middle of the plate (60 wells) were used for cell culture to avoid evaporation. Multiple microplates of identical cell cultures were prepared for temporal evaluation of the cell culture. These plates were subjected to the chemical assay sequentially at 48, 96, 144, and 168 h. 10 μl of CCK-8 (Dojin) was added to the cell culture and incubated at 37°C for one hour, according to the protocol. Finally, 20 μl of 0.1 M HCl was added to arrest the reaction, and the absorbance at 450 nm was measured with a plate reader (Epoch 2, BioTek). A450 was used as the relative cell concentration for machine learning.

### Data processing

Absorbance reads of the chemical assay were exported from the plate reader and processed with Python, as described previously^30^. The mean A450 of biological replications was calculated using “mean” in the “numpy” library. The actual A450 of the cell culture was calculated by subtracting the mean A450 of the medium. The datasets obtained in the present study were summarized in Table S1 and Table S2.

### Machine learning

The algorithm of gradient-boosting decision tree (GBDT) was used in machine learning (ML), which was performed with Python, as described previously^30^. The medium components and A450 of cell culture were employed as the explanatory and objective variables, respectively. The “GradientBoostingRegressor” of the “ensemble” module of the “scikit-learn” library was used to construct the ML model. Five-fold cross-validation and grid search were performed to search for hyperparameters, which used “GridSearchCV” in the “model_selection” module of the “scikit-learn” library. The hyperparameters were searched for “learning_rate” from 0.001 to 0.5 in increments of 0.005, “max_depth” from 2 to 5 in increments of 1, and n_estimators fixed at 300. The other hyperparameters were used by default. In addition, the “feature_importances_” attributed to the GBDT model constructed in the outer five-fold cross-validation was used. Five replications were performed, and the mean values were used as the final evaluation.

### Active learning for medium optimization

The active learning of either regular or time-saving mode was performed with a supercomputer, the Cygnus system (NEC LX 124Rh-4G). According to the ML model constructed with the initial training dataset, approximately 10 million candidate medium combinations were obtained by altering the concentrations of the medium components with four to five variations. By inputting the 10 million candidate media into the ML model, the relative cell culture, represented by A450, was predicted. The top 18 or 19 medium combinations of high A450 were selected and subjected to experimental verification. The experimental results were added to the training dataset for the following learning and prediction. The learning, prediction, and validation were performed repeatedly to achieve the improved cell culture, that is, as high A450 as possible. In addition, the experimental results were used to evaluate the prediction accuracy of the ML models.

### Evaluation of the ML models

Five-fold nested cross-validation was performed to calculate the prediction accuracy of the ML models, as described previously^30^. The five scores computed in the outer 5-fold cross-validation were used for prediction accuracy. Three metrics were employed to evaluate the prediction accuracy: mean absolute error (MAE), coefficient of determination (R^2^), and root mean square error (RMSE). MAE and R^2^ were calculated using the “mean_absolute_error” and “ r2_score “ in the “metrics” module of the “scikit-learn” library. RMSE was calculated using “mean_squared_error” in the “metrics” module of the “scikit-learn”. The square root of the MSE was calculated using “sqtr” in the “numpy” library.

## Acknowledgments

This work was supported by the JSPS KAKENHI Grant-in-Aid for Challenging Exploratory Research (21K19815) and partially by Grant-in-Aid for Scientific Research (B) (19H03215).

## Author contributions

BWY conceived the research; TH and YO performed the experiments; TH and BWY analyzed the data and wrote the paper; and all the authors approved the paper.

## Competing interests

The authors declare no competing interests.

## Data availability

All data generated or analyzed during this study are included in this published article and its supplementary information files Tables S1 and S2.

## Notes

### Competing Interest Statement

The authors have declared no competing interest.

