## Supplementary material for "Employing active learning in the optimization of culture medium for mammalian cells": Figure S1∼S9

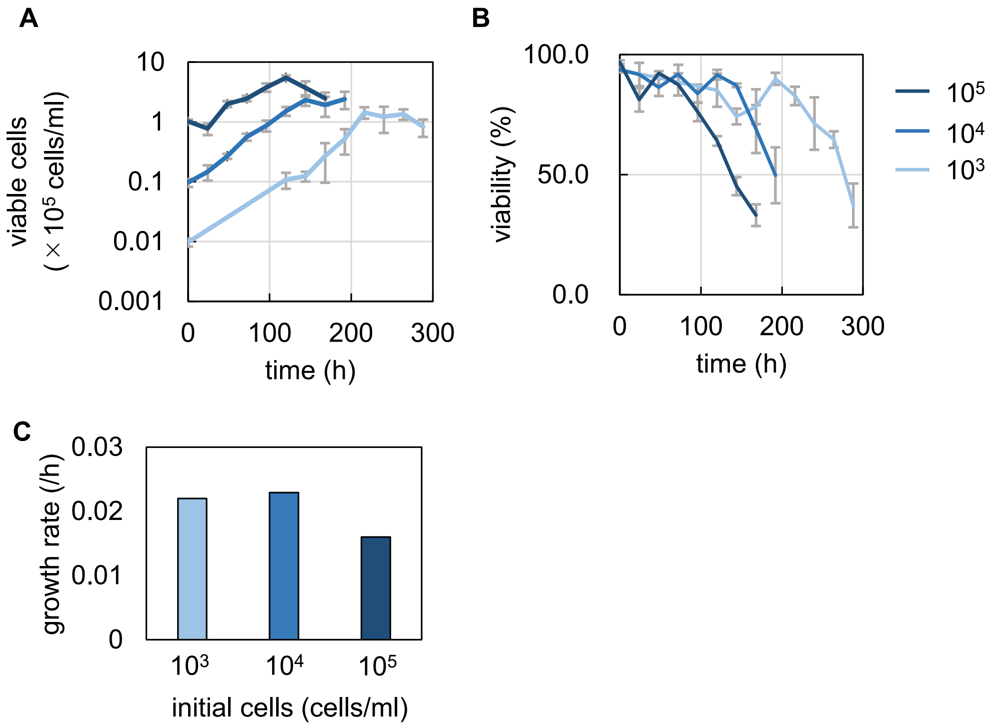


**Figure S1 Evaluation of initial cell concentration to cell culture. A.** Temporal changes of cell culture. **B.** Temporal changes in cell viability. **C.** Cell growth rates evaluated by exponential approximation. Color gradation indicates the variation of initial cell concentration, i.e., $10^3$, $10^4$, $10^5$ cells/ml. Standard errors of biological replications are indicated.

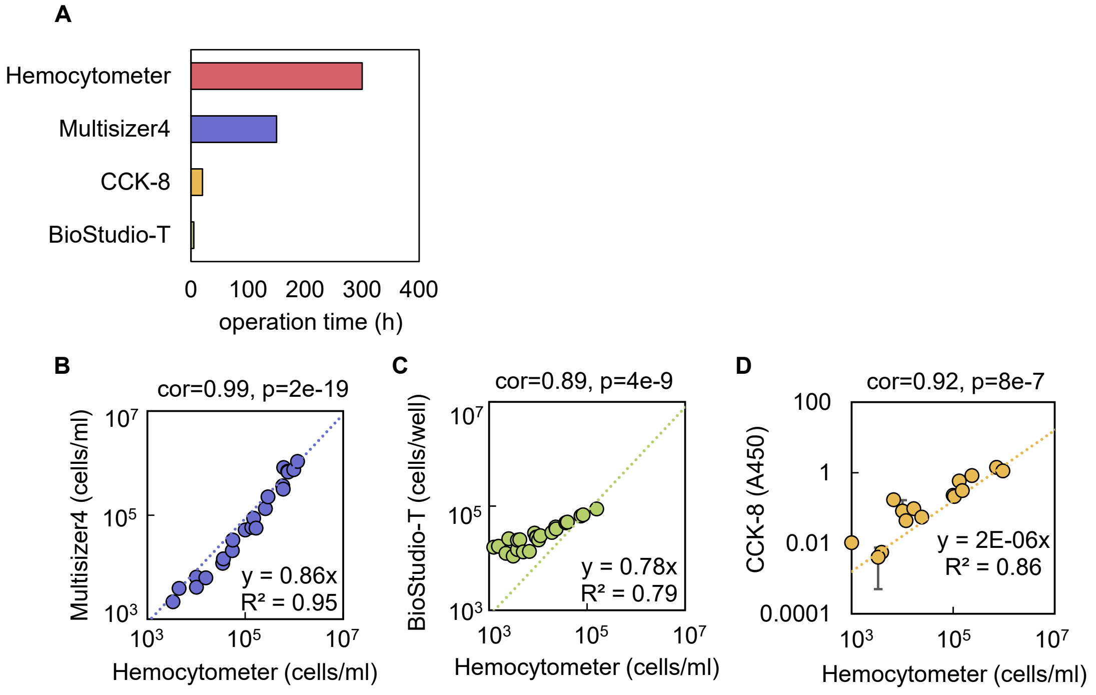


**Figure S2 Various methods for cell counting. A.** Time required to evaluate the cell culture in a 96-well microplate. **B.** Comparison of the cell counting using Multisizer4 and Hemocytometer. **C.** Comparison of the cell counting using BioStudio-T and Hemocytometer. **D.** Comparison of the cell counting using CCK-8 and Hemocytometer. Spearman correlation coefficients and p-values are indicated. The dotted lines represent the linear regression shown with the equations and $R^2$.

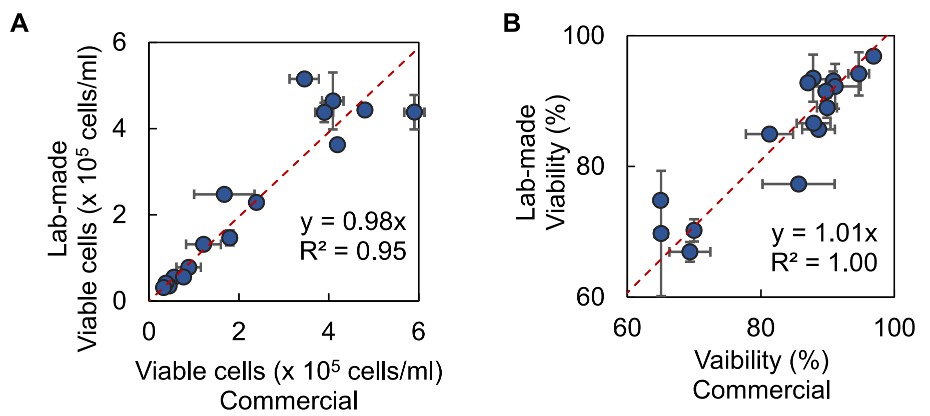


**Figure S3 Comparison of the commercially purchased and lab-made EMEM media.** The number of viable cells (**A**) and viability (**B**) are shown. HeLa cells cultured in 24-well plates for various times (24 ~ 336 h) are shown.

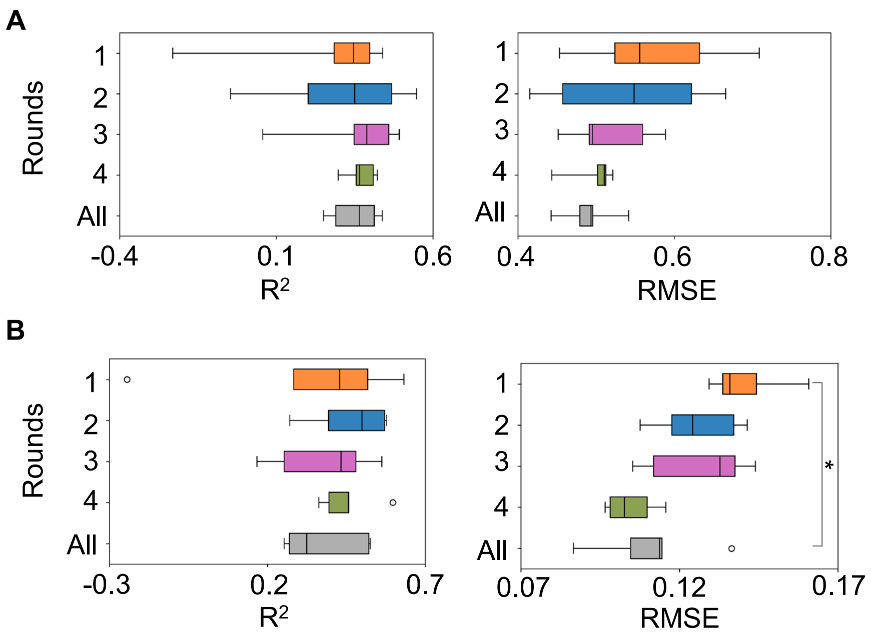


**Figure S4 Prediction accuracy of the ML models. A.** Boxplots of the prediction accuracy in the regular mode. **B.** Boxplots of the prediction accuracy in the time-saving mode. The Left and right panels indicate the metrics of $R^2$ and RMSE, respectively. "All" indicates the accuracy evaluated using the entire dataset from the initial to Round 4. Asterisks indicate statistical significance by Mann-Whitney's U test ($p < 0.05$).

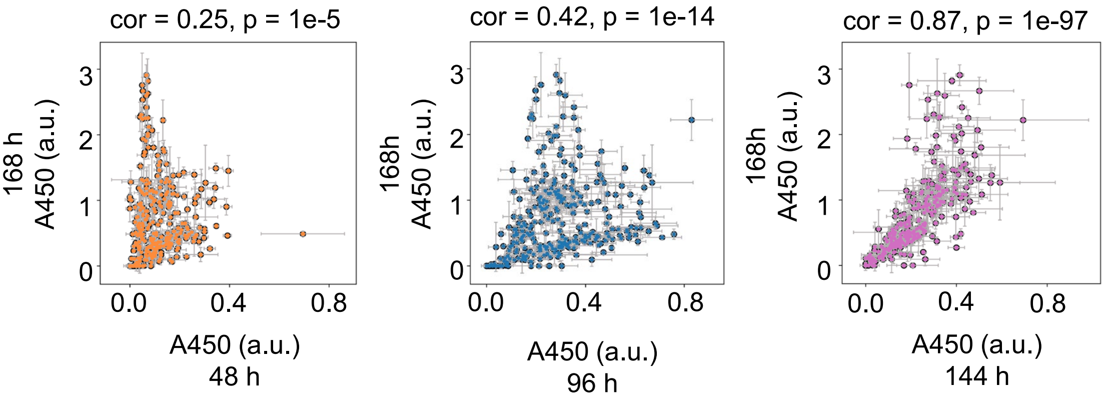


**Figure S5 Relationship of the cell culture at various time points.** Correlations of the cell culture at 168 h to those at 44, 96, and 144 h are shown in orange, blue, and purple, respectively. The cell culture is evaluated as A450. Spearman correlation coefficients and p-values are shown. Standard errors of biological replications are indicated.

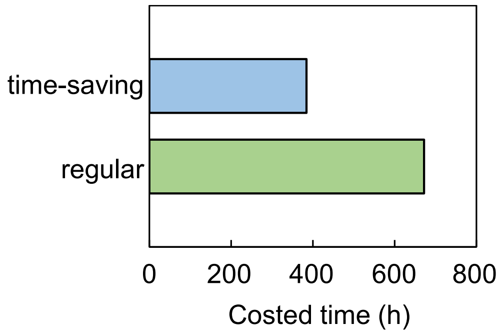


**Figure S6 Time cost for active learning.** The total time cost for the four rounds of active learning was summed.

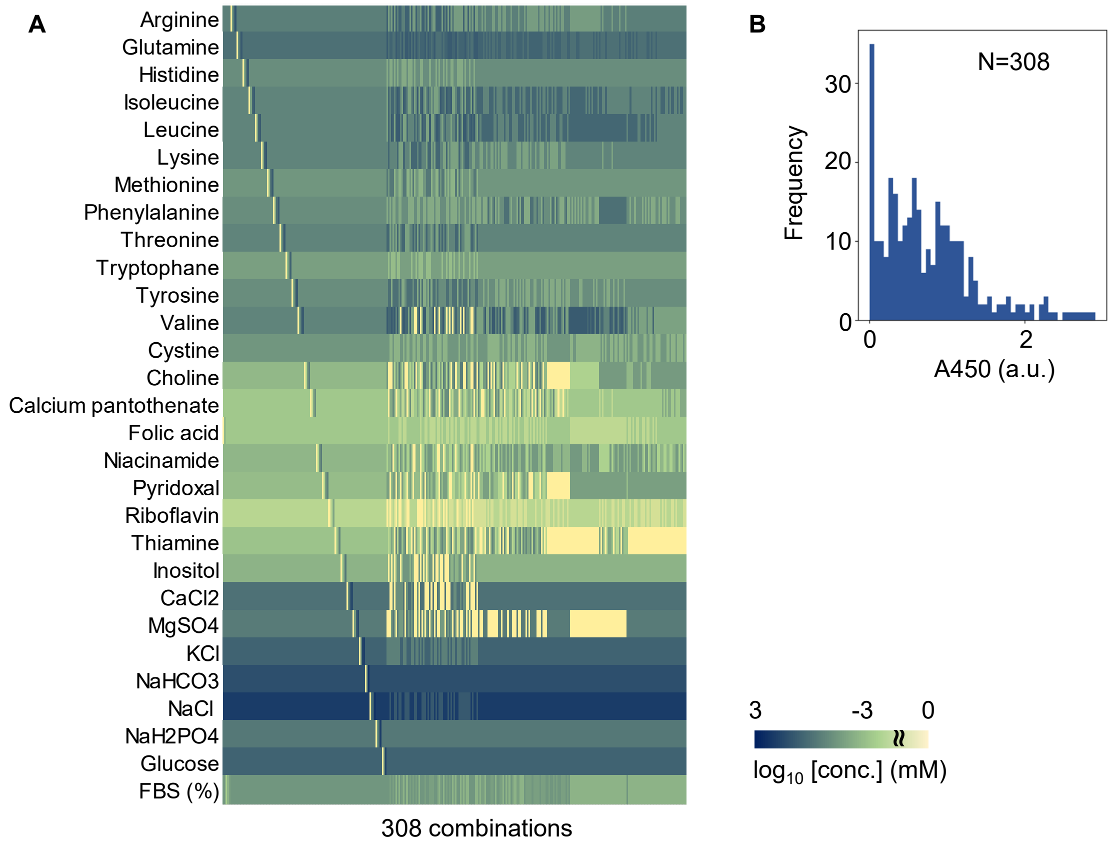


**Figure S7 Medium combinations used in the regular mode. A.** Medium combinations used in active learning. A total of 308 medium combinations are shown in the heatmap. Concentrations are shown on a logarithmic scale. **B.** Distribution of A450 of the cell culture at 168 h. The number of medium combinations is indicated.

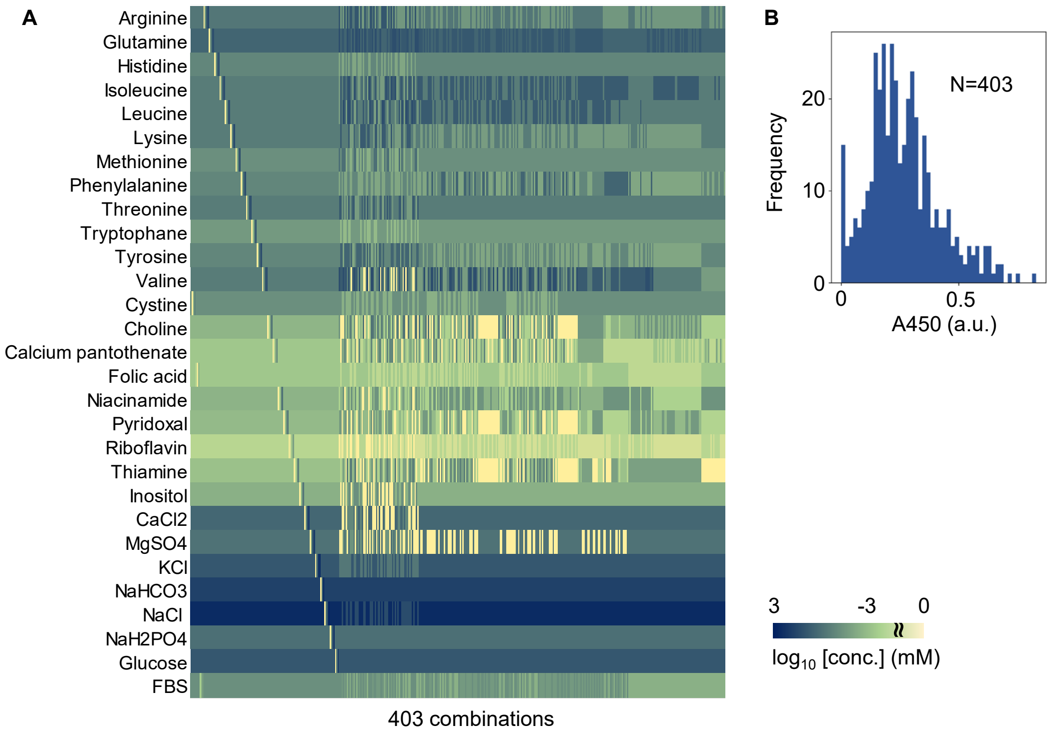


**Figure S8 Medium combinations used in the time-saving mode. A.** Medium combinations used in active learning. A total of 403 medium combinations are shown in the heatmap. Concentrations are shown on a logarithmic scale. **B.** Distribution of A450 of the cell culture at 96 h. The number of medium combinations is indicated.

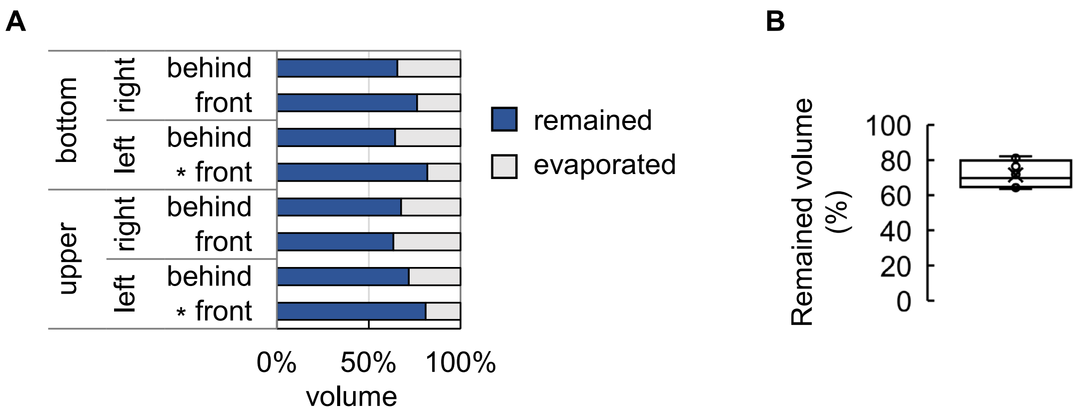


**Figure S9 Evaporation effect during cell culture. A.** Locational dependent evaporation effect. The microtubes containing 1 ml of water were placed at eight different locations in the incubator at 37°C for one week, as indicated. The evaporation effect was evaluated according to the weight of water remaining in the tube. Asterisks indicate significant evaporation, which was evaluated by Mann-Whitney's U-test ($p<0.05$). **B.** Boxplot of the evaporation effect at the eight locations.
